# GroundAnnot: a closed-vocabulary contract for grounding LLM gene-set annotation in live enrichment backends

**DOI:** 10.64898/2026.09.27.754741

**Authors:** Maano Malima

## Abstract

**Motivation:** LLM agents increasingly draft functional interpretations of gene lists, but can cite Gene Ontology (GO) terms that no current enrichment backend returned for that list, and can pair real GO accessions with fabricated labels.

**Results:** We present GroundAnnot, a client for PANTHER, Enrichr, and g:Profiler that returns a closed vocabulary of GO term IDs and their backend labels, and enforces two contracts: Contract A (no ID or label outside the backend payload) and Contract B (no enrichment claim outside the backend’s significant set). Across three open-weight models (Qwen2.5-7B local; Qwen3.8-27B and GPT-OSS-120B via Groq) on six curated disease gene lists, mean valid-enriched rates were 15.3%, 20.1%, and 35.1%. All unsupported IDs were real GO terms classified by QuickGO as wrong-biology, obsolete, or wrong-branch (zero fabricated accessions). A distinct failure mode emerged: for the Alzheimer’s gene list, the local 7B model paired every one of its ten stable picks with a label that does not match the current GO term for that accession (10/10 mismatches), producing a coherent synaptic-signalling narrative that the IDs do not support. Mean overlap with the backends’ FDR top-10 union was 0.33–0.67/10. On 50 MSigDB Hallmark gene sets, the strongest model named the defining pathway in 21/44 (47.7%) directly named cases. When an enriched shortlist was placed in the prompt, all three models complied fully (60/60 for each model; 180/180 overall), showing that a bounded vocabulary removes these output classes when the downstream agent is required to use it.

**Availability and Implementation:** GroundAnnot is implemented in Python 3.9+ and is freely available on GitHub at https://github.com/Maano-gen/groundannot under the MIT license, together with the six disease gene lists, 50 Hallmark gene lists, all 215 disease-list and 150 Hallmark runs, two dated backend snapshots, and the analysis scripts.

**Supplementary Information:** Supplementary data are available online.

## 1 Introduction

After RNA-seq, a CRISPR screen, or variant-to-gene analysis, biologists need functional labels for a gene list: GO processes, molecular functions, pathways. PANTHER (Thomas et al., 2022; Mi et al., 2021), Enrichr (Chen et al., 2013; Kuleshov et al., 2016), and g:Profiler (Raudvere et al., 2019) perform this task with live curated data and expose programmatic access. Separately, LLM agents increasingly draft interpretations in natural language. The failure mode of this second layer is documented. Joachimiak et al. (2023) showed that GPT-based gene-set summarisation can emit plausible GO terms that are not recovered by standard statistical enrichment, and concluded that LLM methods are unsuitable as a replacement for term-enrichment analysis. Hu et al. (2025) evaluated large language models for gene-set function discovery and found stronger recovery of curated set names than of omics-derived clusters. GeneAgent (Wang et al., 2025) addresses grounding by generating terms and then verifying them against biological databases.

GroundAnnot takes a different approach. Rather than verify after generation, it constrains generation prospectively (Fig. 1). The tool queries the three backends once, writes two bounded vocabularies to disk (allowed_term_ids.json: every ID any backend returned, with its backend label; enriched_term_ids.json: the subset each backend flags as significant at FDR < 0.05 or the backend’s equivalent default), and constrains any downstream LLM to select from those sets. The model is a narrator, not an annotation authority.

**Figure 1.**
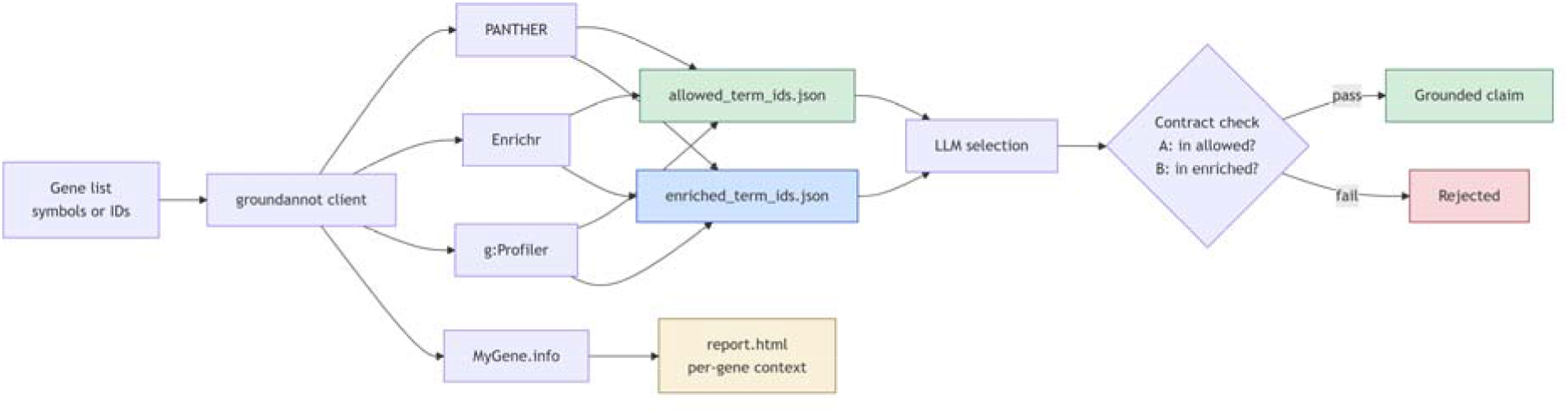
The GroundAnnot pipeline. A gene list is sent to three live enrichment backends (PANTHER, Enrichr, g:Profiler); the client writes two bounded vocabularies (allowed_term_ids.json, enriched_term_ids.json) and a per-gene context report. A downstream LLM selects terms from those vocabularies. The contract check accepts claims whose IDs and labels match the backend payload and rejects the rest.

We report three findings. First, unconstrained open-weight LLMs produce valid enriched terms at mean rates of 15–35% on these lists, and their picks do not track the backends’ own significance ranking. Second, the same models pair real GO accessions with fabricated labels; for one Alzheimer’s list, all ten picks carried incorrect labels. Third, when the enriched shortlist is placed in the prompt, all three models comply at 100%. The failure is not model refusal; it is the absence of a bounded vocabulary.

## 2 System and Methods

### 2.1 Backends

GroundAnnot queries three enrichment backends: PANTHER (endpoints: POST /services/oai/pantherdb/geneinfo and /enrich/overrep) (Thomas et al., 2022; Mi et al., 2021), Enrichr (POST /addList, GET /enrich) (Chen et al., 2013; Kuleshov et al., 2016), and g:Profiler (POST /api/gost/profile/) (Raudvere et al., 2019). MyGene.info (Xin et al., 2016) provides per-gene context (name, summary, GO terms, pathways, InterPro) and is not used for enrichment claims. DAVID was excluded after DNS failures on legacy hosts and 404 responses from the REST paths probed at the time of this study (2026-09-23). Other endpoint paths may exist; we did not exhaustively probe DAVID’s current web services.

### 2.2 Contracts

- Contract A: every GO ID cited by a downstream agent must appear in allowed_term_ids.json, and the label the agent displays must match the backend label for that ID.
- Contract B: every claim of enrichment must cite an ID in enriched_term_ids.json.

Both contracts follow the backends’ own default behaviour: PANTHER displays only FDR < 0.05 unless “display all results” is selected; g:Profiler defaults user_threshold = 0.05; Enrichr reports Benjamini–Hochberg-adjusted p-values, with common practice at ≤ 0.05. For g:Profiler, enriched terms carry g:SCS-adjusted p-values; the backend does not emit a separate FDR column. Reporting follows the minimum standards of Wijesooriya et al. (2022). In practice allowed_term_ids.json is large (roughly 13,000-14,000 IDs per disease list, pooled across the three backends), so Contract A alone is a weak filter on its own; most of the discriminative work is done by Contract B (the enriched subset) together with the label check in §2.6, and readers should not read “allowed” as “enriched.”

### 2.3 Test gene lists

We used six curated disease panels comprising 121 genes in total: Alzheimer’s disease (15; familial AD), hereditary cancer (28; cancer-related subset of ACMG SF v3.2), hypertrophic cardiomyopathy (20; sarcomere literature), Parkinson’s disease (16; monogenic PD), primary immunodeficiency (24; IUIS core), and type 2 diabetes (18; GWAS consensus). Fifty MSigDB Hallmark gene sets (Liberzon et al., 2015) were taken from the Enrichr library MSigDB_Hallmark_2020 and used as a second test set whose defining biology is encoded in the set name.

### 2.4 Models and runs

We selected three open-weight models based on the availability of inspectable weights or public inference endpoints: Qwen2.5-7B-Instruct (Q4_K_M GGUF; local llama.cpp; temperature 0; seed 42), Qwen3.8-27B (Groq model ID qwen/qwen3.8-27b), and GPT-OSS-120B (Groq model ID openai/gpt-oss-120b; reasoning effort set to low). Qwen3.8-27B was accessed via the Groq API as a preview model (model ID qwen/qwen3.8-27b; Groq, 2026). The model is a 27B multimodal open-weight model from the Qwen series. The local model produced bit-identical output across 36 disease-list runs (SD = 0.00 on all scoring metrics). The two hosted models show provider-side nondeterminism; 92 and 87 disease-list runs were collected, respectively. On the 50 Hallmark sets, one run per model was collected for theme-recovery scoring. Two backend snapshots collected on consecutive days (2026-09-24 and 2026-09-25) were pulled for the same six disease lists.

#### Unconstrained prompt (verbatim)

System: “You are a bioinformatician. Answer concisely and only in the requested format.” User: “List exactly 10 Gene Ontology biological process terms that are enriched in this gene list. Gene list: {genes}. Output format: one per line, no preamble, no numbering: GO:XXXXXXX term name.” Temperature was set to 0 for all three models; Groq requests for GPT-OSS-120B additionally set reasoning_effort = low.

### 2.5 Scoring

For each unconstrained LLM run, every emitted GO ID was classified as valid enriched (present in enriched_*), allowed-not-enriched (present in allowed_* but not enriched_*), or unsupported (absent from every backend payload for that list). Every unsupported ID was checked live against QuickGO and labelled WRONG_BIOLOGY, OBSOLETE, WRONG_BRANCH, NOT_IN_GO, or UNKNOWN. Rates below are means across the 215 disease-list runs unless noted. Hallmark scoring used the 150 hallmark runs.

### 2.6 Label agreement

For each pick, the label the model produced was compared against the live QuickGO label for the corresponding accession (normalised by lowercasing and stripping non-alphanumeric characters). A mismatch was recorded when the normalised strings differed. This measures label correctness independently of ID correctness.

### 2.7 Bounded-prompt condition

The bounded prompt included the union of the top-50 terms by lowest FDR from each backend (deduplicated; shortlist size ≤ 150). The prompt instructed the model to select 10 terms from this list.

#### Bounded prompt (verbatim)

System: “You are a bioinformatician. You must answer only using the provided list.” User: “You are given a gene list and a shortlist of Gene Ontology biological-process terms that are statistically enriched in that gene list. Gene list: {genes}. Enriched GO terms (pick ONLY from this list, use the exact GO ID): {shortlist}. Select exactly 10 of the terms above that best describe the biology of the gene list. Output one per line, no preamble, no numbering: GO:XXXXXXX term name.”

## 3 Results

### 3.1 Unconstrained LLM output

Table 1 shows mean valid-enriched rates on six disease lists.

**Table 1.** Mean valid-enriched rates on six disease lists. CI bounds are from 1,000 bootstrap resamples of runs, reported by stats/bootstrap_ci.csv. The local model produced bit-identical output across all 36 runs, so its interval collapses to the point estimate. Rates are multi-run means; the 18/32/9, 9/38/12, and 8/31/21 compositions quoted in the text are from a single 6-list output per model, not pooled across runs.

| Model | Disease-list runs | Mean valid-enriched rate | Bootstrap 95% CI | Mean unsupported rate (Contract A) |
| --- | --- | --- | --- | --- |
| Qwen2.5-7B-Instruct (local) | 36 | 15.25% | [15.25%, 15.25%] | 31% |
| Qwen3.8-27B (Groq) | 92 | 20.14% | [19.81%, 20.45%] | 15% |
| GPT-OSS-120B (Groq) | 87 | 35.05% | [34.64%, 35.50%] | 13% |

**Table 2.** Hallmark direct recovery per model.

| Model | Direct recovery |
| --- | --- |
| GPT-OSS-120B | 21/44 (47.7%) |
| Qwen3.8-27B | 8/44 (18.2%) |
| Qwen2.5-7B | 6/44 (13.6%) |

A single 6-list output from each model contained 59 or 60 emitted IDs. In that snapshot, composition was 18/32/9 (unsupported / allowed-not-enriched / valid enriched) for Qwen2.5-7B, 9/38/12 for Qwen3.8-27B, and 8/31/21 for GPT-OSS-120B. Across all runs, QuickGO returned NOT_IN_GO = 0. Unsupported IDs were real GO terms used as wrong-biology, obsolete, or wrong-branch labels.

### 3.2 Overlap with FDR ranking

For each disease, the top-10 terms by lowest FDR from each backend were pooled (union ≤ 30) and compared with each run’s 10 picks. Mean overlap: 0.67/10 (GPT-OSS-120B), 0.33/10 (Qwen3.8-27B), 0.33/10 (Qwen2.5-7B).

As an informal order-of-magnitude check, if 10 terms are drawn uniformly from an enriched pool of ∼400 IDs, expected overlap with a 30-term FDR union is 10 × 30/400 = 0.75. That is an upper bound on this null: the union is often smaller than 30, and the enriched-pool size varies by disease (tens to hundreds of terms). Observed means sit at or below that bound and do not follow the backends’ significance ranking. Under a full-GO draw (∼20,000 terms) the same null is ∼0.005–0.015/10. This calculation is not a formal inferential null model. The models are closer to “any enriched term” than to “top by FDR,” and closer to neither than to a calibrated ranker.

### 3.3 Hallmark theme recovery

On the 44 Hallmarks whose defining biology is named in the set title (DNA_REPAIR, APOPTOSIS, HYPOXIA, and similar), a pick counted as a hit if at least one of the ten labels contained a keyword for that name (lexical match, not expert review):

The six indirect Hallmarks, MYC_TARGETS_V1, MYC_TARGETS_V2, E2F_TARGETS, KRAS_SIGNALING_UP, KRAS_SIGNALING_DN, and G2M_CHECKPOINT, had 0/6 recovery for every model. That split is expected under lexical scoring: the right terms there name downstream processes, not the regulator (Fig. 2).

**Figure 2.**
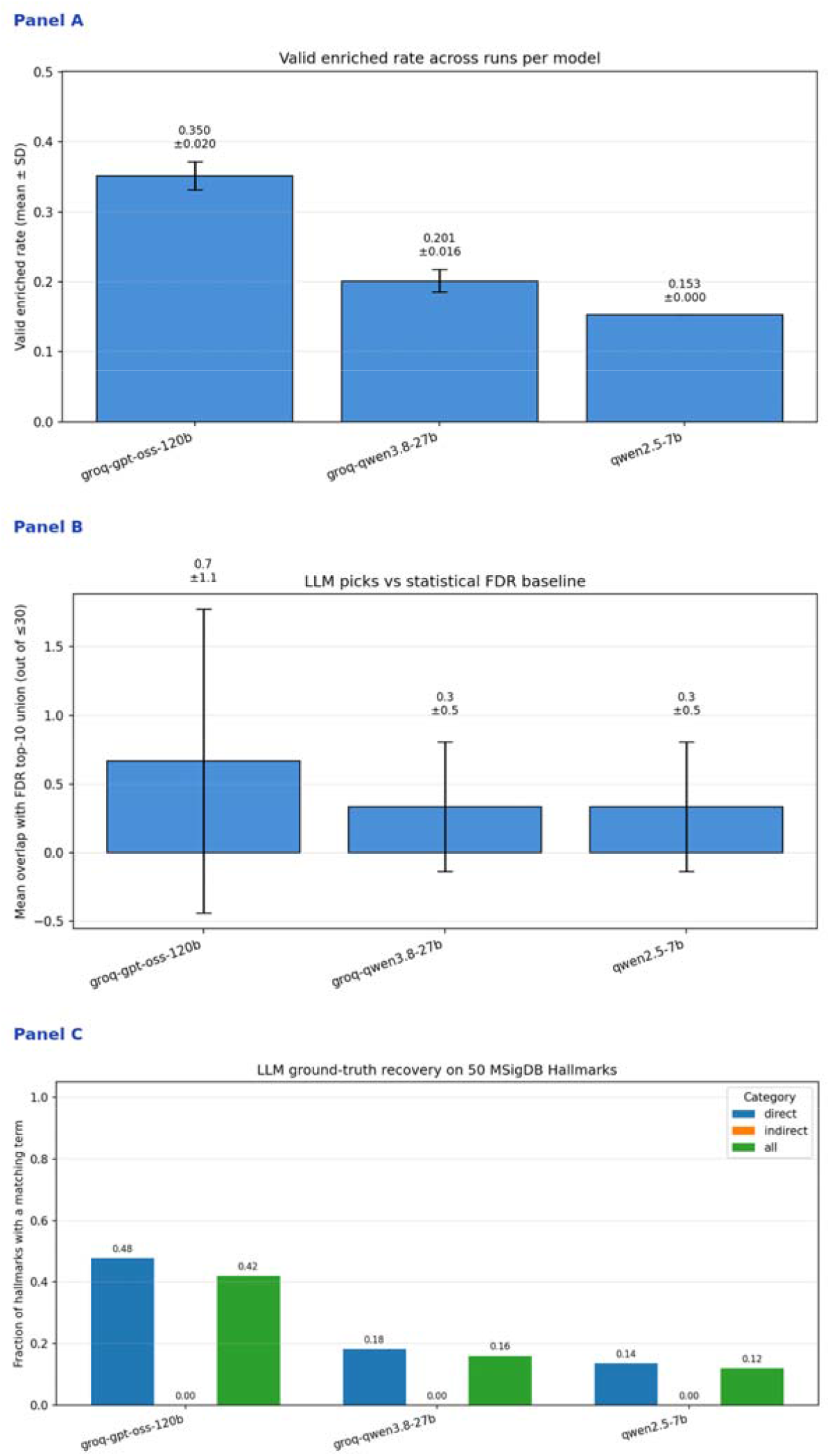
Summary results. Panel A: mean valid-enriched rates per model; all 36 local-model runs produced identical output, and the 95% bootstrap CI on the valid-enriched rate collapses to the point estimate 15.25%. Panel B: mean overlap with the FDR top-10 union per model. Panel C: MSigDB Hallmark direct recovery per model.

### 3.4 Reproducibility without correctness

The local Qwen2.5-7B was bit-identical across 36 runs (SD = 0.00 on all metrics). Its stable top-10 picks for the Alzheimer’s list were animal-behaviour GO terms, including fly-neuroscience classics such as anesthesia-resistant memory (GO:0007615) and courtship behaviour (GO:0007619). Those ten IDs were present in every local run, had zero overlap with the backends’ FDR top-10 union for that list, and carried mixed ID-level contract status: some valid enriched, some allowed-not-enriched, some unsupported b any backend. All ten, regardless of ID-level status, carried a label that did not match the backend’s own label for that accession (10/10 mismatches; see §3.5). A deterministic model can return the same biologically inappropriate answer in every run.

### 3.5 Fabricated labels for real IDs

Every one of the ten stable picks carried a label that does not match the live QuickGO label for that accession (10/10 mismatches; Fig. 3). The model’s claimed labels, “synaptic transmission,” “postsynaptic membrane,” “presynaptic membrane,” “neurotransmitter secretion,” and “synaptic vesicle exocytosis”, were internally coherent and biologically plausible for an Alzheimer’s gene list. The ten accessions, however, all fall in the range GO:0007601–GO:0007632, with current QuickGO labels including visual perception, learning, mating behaviour, courtship behaviour, rhythmic behaviour, ultradian rhythm, locomotory behaviour, adult walking behaviour, and visual behaviour. The model produced a coherent synaptic-signalling narrative that the IDs do not support.

**Figure 3.**
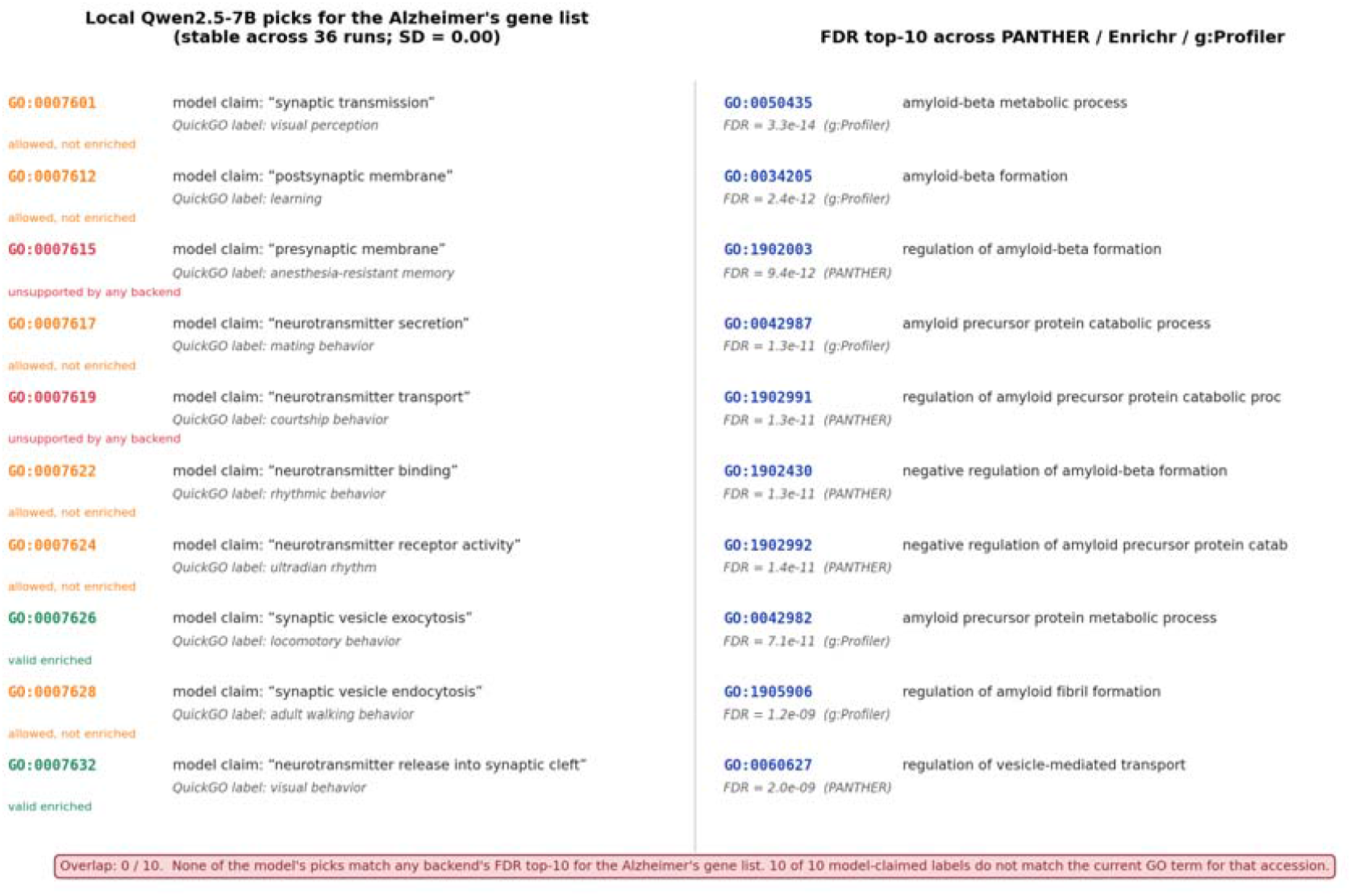
Alzheimer’s gene list. Left: the local Qwen2.5-7B model’s ten stable picks; all 36 runs produced identical output. Each entry shows the model-claimed label above the live QuickGO label for the same accession; categories below each entry show contract status (green = valid enriched, orange = allowed but not enriched, red = unsupported by any backend). Right: the FDR top-10 union across the three backends, with the backend and FDR value shown for each. None of the model’s picks match the FDR top-10. All ten model-claimed labels differ from the QuickGO label for the accession.

This is a distinct failure mode from “fabricated accessions.” A downstream agent that trusts the displayed label without checking the ID-to-label mapping would not detect it. Contract A, as enforced by GroundAnnot, matches both ID and backend label, so this class of error is also prevented.

### 3.6 Bounded condition

When the enriched shortlist was placed in the prompt and the model was asked to select from it, all three models complied: 60/60 picks in-list for each model across the six diseases (180/180 overall; in-list rate 1.000).

Applying the same membership filter after the fact to unconstrained runs (retain only picks present in enriched_*) left mean surviving picks of 1.50/10 (Qwen2.5-7B), 1.86/10 (Qwen3.8-27B), and 3.49/10 (GPT-OSS-120B). Zero picks survived in 33%, 17%, and 17% of those runs, respectively. These are means of per-run coverage (kept ÷ emitted). They differ slightly from Table 3 because the hosted models sometimes emitted fewer than 10 IDs.

**Table 3.**
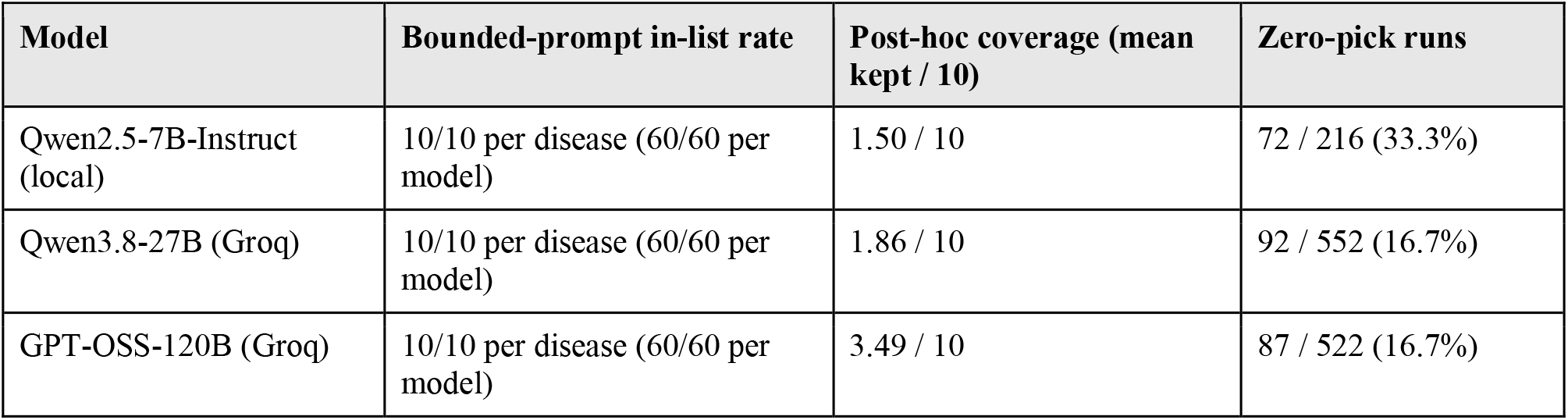
Bounded-prompt compliance versus post-hoc coverage of unconstrained picks. Post-hoc coverage is the mean per-run fraction retained after filtering against enriched_term_ids.json; hosted models occasionally emitted fewer than 10 IDs, so these values differ slightly from the aggregate rates in Table 1. Zero-pick denominators are per-disease run rows (six diseases × model runs). Each model was tested once per disease in the bounded condition (10/10 in-list picks per disease), giving 60/60 in-list picks per model and 180/180 overall.

| Model | Bounded-prompt in-list rate | Post-hoc coverage (mean kept / 10) | Zero-pick runs |
| --- | --- | --- | --- |
| Qwen2.5-7B-Instruct (local) | 10/10 per disease (60/60 per model) | 1.50 / 10 | 72 / 216 (33.3%) |
| Qwen3.8-27B (Groq) | 10/10 per disease (60/60 per model) | 1.86 / 10 | 92 / 552 (16.7%) |
| GPT-OSS-120B (Groq) | 10/10 per disease (60/60 per model) | 3.49 / 10 | 87 / 522 (16.7%) |

The bounded-prompt result shows compliance with an in-list instruction. It does not show that the ten items chosen from the shortlist were the best ten.

### 3.7 Backend drift

Two backend snapshots pulled on consecutive days changed valid-enriched rates by 0–3.3 percentage points for the same model, prompt, and gene list (one observation per cell). A larger same-day swing of about 20 percentage points was seen once during an active backend update. These anecdotal observations support shipping dated snapshots, but the two-day comparison is too small to estimate variance.

## 4 Discussion

GroundAnnot is a small client for three live enrichment backends. It returns two bounded vocabularies and a portable HTML report. The contribution is architectural, not algorithmic. If a downstream agent is actually bound to the JSON, the design removes three failure classes before the model speaks: IDs that no backend returned, enrichment claims that no backend flagged as significant, and IDs paired with labels that do not match the backend’s own label for that accession.

Three empirical points support using that constraint. First, unconstrained open-weight models produced valid enriched terms at 15–35% on these lists, below what an FDR sort scores on the same metric by construction. Second, their picks did not track the backends’ significance ranking (mean FDR-top-10 overlap 0.33–0.67 out of 10). Third, when the enriched shortlist was placed in the prompt, all three models stayed in-list at 60/60 for each model (180/180 overall). The problem in the unconstrained setting was not refusal; it was the absence of a closed vocabulary.

The bit-identical local 7B result is the sharpest illustration. Running that model once, twice, or 36 times yields the same biologically unsupported shortlist, with the same ten accessions, all paired with labels that do not correspond to those accessions. Reproducibility measured by resampling is not correctness, and ID-level validity is not label-level validity.

The label-checking requirement was not part of the original design. It emerged from the benchmark. GroundAnnot’s allowed_* and enriched_* files store both GO IDs and backend labels, so a downstream agent can check that the ID-to-label mapping the LLM claims matches the backend. Removing the label check would leave Contract A satisfiable by a model that uses real IDs with fabricated labels, as the local 7B demonstrated at 10/10.

The work is close in spirit to audit-gated claim systems (Furudate and Takahashi, 2026), which require an LLM’s claims to point back at a machine-readable evidence table. GroundAnnot uses a different mechanism: the vocabulary, not a post-hoc audit, is the constraint. We did not compare the two mechanisms head-to-head.

Unconstrained open-weight models often emit real GO accessions that the live backends did not call enriched, and they can attach the wrong official name to a real accession. A two-file, ID-plus-label contract removes those classes of error if the agent is required to use it.

## 5 Limitations

- No frontier or tool-using model was tested. GPT-4, Claude, and Gemini were excluded because weights and inference stacks are not publicly verifiable. GeneAgent-style generate-then-verify was not compared. Reported rates are for unconstrained open-weight models.
- Hallmark scoring is lexical, not semantic. Theme recovery asks whether one of ten labels contains a keyword from the Hallmark name. It is not expert agreement. The direct/indirect split is an acknowledgement of that limit.
- Backend drift was measured at N = 1 per snapshot day. The 0–3.3 point consecutive-day difference is not a variance estimate. The single ∼20-point same-day swing during an update is the stronger drift observation.
- Disease lists are small (15–28 genes). Generic enrichment terms appear more readily on small lists. Raw valid-enriched rates should be read with that in mind.
- The bounded-prompt test measures compliance, not ranking quality. Selecting 10 items from a shortlist of up to 150 shows that the models follow the contract. It does not show that they selected the best 10.
- The label-mismatch result is detailed for one disease list. The 10/10 mismatch figure is reported for the Alzheimer’s list only. A full label-agreement audit across all 215 runs is future work.
- DAVID was not included because the legacy hosts and REST paths probed on 2026-09-23 failed. Other current endpoint paths may exist, and the available DAVID web services were not exhaustively tested. The multi-backend claim is limited to PANTHER, Enrichr, and g:Profiler.
- Local determinism covers one model. Bit-identical output was verified for Qwen2.5-7B-Instruct (Q4_K_M GGUF) via llama.cpp. It was not verified for the hosted models.
- All benchmark numbers are author-computed from the deposited run JSONs and have not been independently reproduced.
- Table 1 rates are multi-run means; the 59/60-ID breakdown is a single output composition. Do not read 18 unsupported IDs as a count pooled over 36 runs.
- ACMG provenance is a subset. The 28-gene hereditary-cancer panel is not the full ACMG SF v3.2 list.

## 6 Availability

Source code, six disease gene lists, 50 Hallmark gene lists, all 215 disease-list runs and 150 Hallmark runs, two dated backend snapshots, and the analysis scripts are available on GitHub at https://github.com/Maano-gen/groundannot under the MIT license. The GroundAnnot package requires Python 3.9+, requests, and jinja2. Analysis scripts additionally require pandas and matplotlib. The optional LLM benchmark requires llama-cpp-python (local) or openai (API) and a Groq API key in .env.

## Acknowledgements

I would like to thank Setshaba Taukobong for the discussion that led to this paper.

## Data Availability

The six curated disease gene lists, 50 MSigDB Hallmark gene lists, all 215 disease-list and 150 Hallmark LLM runs, two dated backend snapshots (2026-09-24 and 2026-09-25), and the analysis scripts underlying this article are available on GitHub at https://github.com/Maano-gen/groundannot under the MIT license.

## References

Chen, E.Y., Tan, C.M., Kou, Y., Duan, Q., Wang, Z., Meirelles, G.V., Clark, N.R., Ma’ayan, A. (2013) Enrichr: interactive and collaborative HTML5 gene list enrichment analysis tool. BMC Bioinformatics, 14, 128.

Furudate, K., Takahashi, K. (2026) LLM-PathwayCurator transforms enrichment terms into audit-gated decision-grade claims. bioRxiv. doi:10.64898/2026.02.18.706381.

Groq. (2026) Supported models. https://console.groq.com/docs/models

Hu, M., Alkhairy, S., Lee, I., Pillich, R.T., Fong, D., Smith, K., Bachelder, R., Ideker, T., Pratt, D. (2025) Evaluation of large language models for discovery of gene set function. Nature Methods, 22, 82–91.

Joachimiak, M.P., Caufield, J.H., Harris, N.L., Kim, H., Mungall, C.J. (2023) Gene set summarization using large language models. arXiv:2305.13338.

Kuleshov, M.V., Jones, M.R., Rouillard, A.D., Fernandez, N.F., Duan, Q., Wang, Z., Koplev, S., Jenkins, S.L., Jagodnik, K.M., Lachmann, A., McDermott, M.G., Monteiro, C.D., Gundersen, G.W., Ma’ayan, A. (2016) Enrichr: a comprehensive gene set enrichment analysis web server 2016 update. Nucleic Acids Research, 44, W90–W97.

Liberzon, A., Birger, C., Thorvaldsdóttir, H., Ghandi, M., Mesirov, J.P., Tamayo, P. (2015) The Molecular Signatures Database (MSigDB) hallmark gene set collection. Cell Systems, 1, 417–425.

Mi, H., Ebert, D., Muruganujan, A., Mills, C., Albou, L.P., Mushayahama, T., Thomas, P.D. (2021) PANTHER version 16: a revised family classification, tree-based classification tool, enhancer regions and extensive API. Nucleic Acids Research, 49, D394–D403.

Raudvere, U., Kolberg, L., Kuzmin, I., Arak, T., Adler, P., Peterson, H., Vilo, J. (2019) g:Profiler: a web server for functional enrichment analysis and conversions of gene lists (2019 update). Nucleic Acids Research, 47, W191–W198.

Qwen. (2026) Qwen3.8-27B model card. Hugging Face. https://huggingface.co/Qwen/Qwen3.8-27B

Thomas, P.D., Ebert, D., Muruganujan, A., Mushayahama, T., Albou, L.P., Mi, H. (2022) PANTHER: Making genome-scale phylogenetics accessible to all. Protein Science, 31, 8–22.

Wang, Z., Jin, Q., Wei, C.-H., Tian, S., Lai, P.-T., Zhu, Q., Day, C.-P., Ross, C., Leaman, R., Lu, Z. (2025) GeneAgent: self-verification language agent for gene-set analysis using domain databases. Nature Methods, 22, 1677–1685.

Wijesooriya, K., Jadaan, S.A., Perera, K.L., Kaur, T., Ziemann, M. (2022) Urgent need for consistent standards in functional enrichment analysis. PLoS Computational Biology, 18, e1009935.

Xin, J., Mark, A., Afrasiabi, C., Tsueng, G., Juchler, M., Gopal, N., Stupp, G.S., Putman, T.E., Ainscough, B.J., Griffith, O.L., Torkamani, A., Whetzel, P.L., Mungall, C.J., Mooney, S.D., Su, A.I., Wu, C. (2016) High-performance web services for querying gene and variant annotation. Genome Biology, 17, 91.

